## Supplementary Information for "*Plasmodium* ARK1 regulates spindle formation during atypical mitosis and forms a divergent chromosomal passenger complex"

### Supplementary Figures

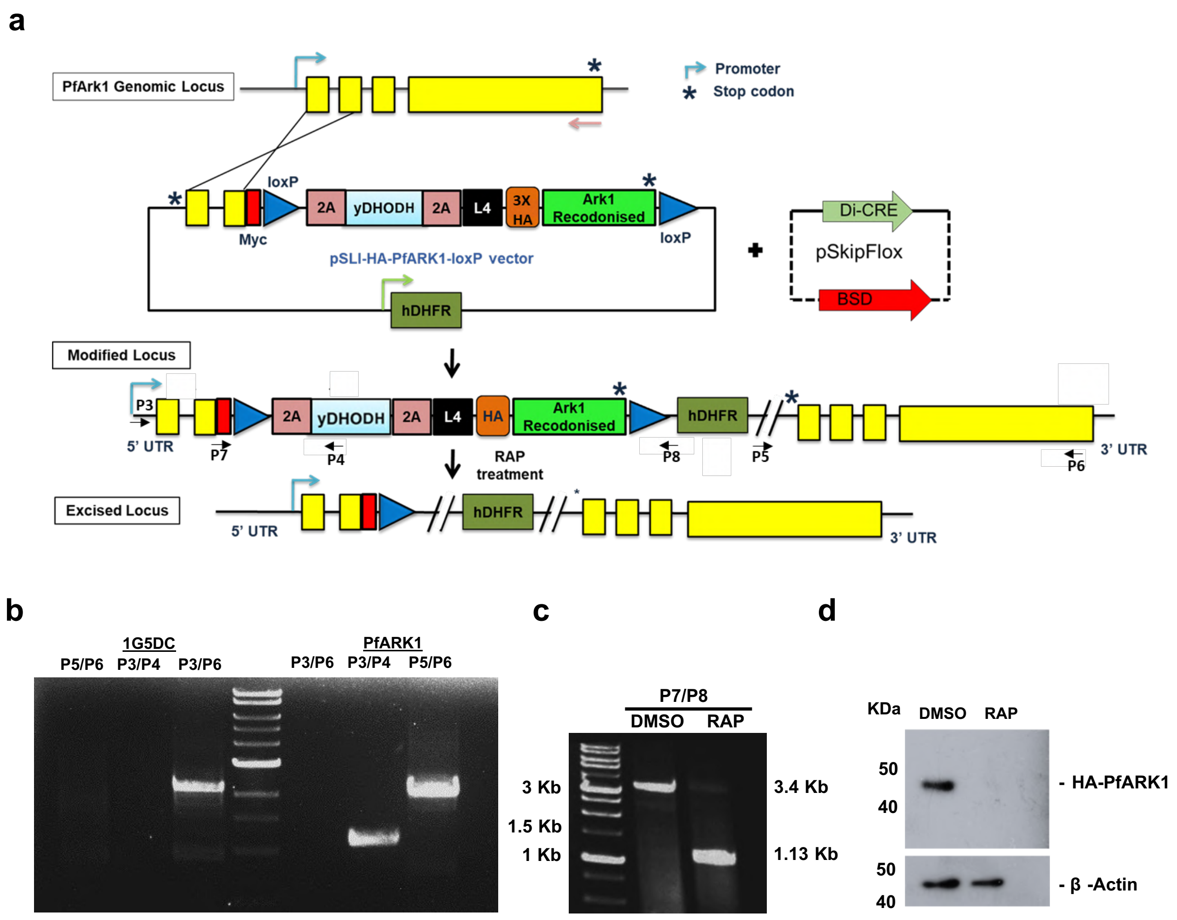


| Locus | Primerset | Expected size |
|---|---|---|
| WT | P3/P6 | 2.2 Kb |
| 5'Integration | P3/P4 | 1.11 Kb |
| 3'Integration | P5/P6 | 1.9 Kb |

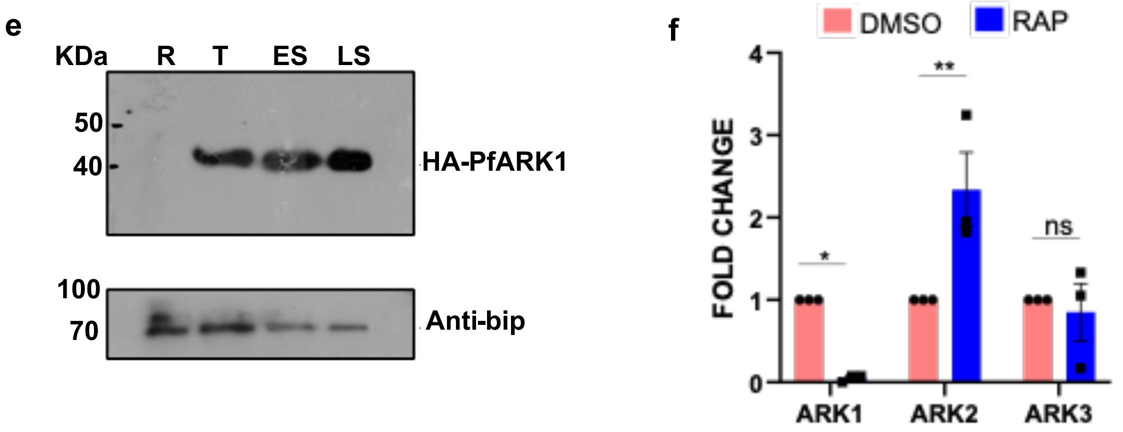

**Supplementary Figure 1. Conditional Knockout of PfARK1 and effect on expression of other PfARKs.**

**a**. Schematic representation of the strategy used to modify the PfARK1 locus using selection-linked integration (SLI)[1, 2]. A homology region corresponding to the 5'-end of PfARK1was cloned in pSLI-N-sandwich-loxP vector. In addition, recodonised PfARK1 with N-terminal 3xHA tag was cloned downstream of yDHODH, which was spanned by 2A skip peptide. As a result, 3xHA tagged ARK1 would be expressed upon recombination. *-stop codon, arrows-location of PCR primers that were used for genotyping and assessing excision. The above-mentioned construct was transfected in 1G5DC parasites followed by pSkipFlox plasmid, which contains a spilt Cre recombinase insert and parasites were treated with WR99210 and DSM1 as described in experimental procedures. **b**. Genotyping of HA-PfArk1-loxP parasites that appeared after drug selection. PCR amplification was carried out using primers targeting the indicated sites in panel a. PCR products of expected size, which is indicated in the table, were amplified and suggested the desired 5'- and 3'-integration whereas the wild-type (WT) locus was absent in the transgenic HA-PfARK1-loxP parasites. The amplicons were sequenced to further confirm the integration at the desired locus. **c**. HA-PfARK1-loxP parasites were treated with DMSO or 250 nM RAP for 24h. Subsequently, genomic DNA was isolated at ~44 hpi and PCR was performed using primer sets P7 and P8. A ~1.13kb product was obtained upon RAP treatment indicating successful excision. **d**. Western blot examination of HA-PfARK1-loxP parasite lysates (~44 hpi, cycle 0) treated with DMSO or RAP and using anti-HA antibody revealed that the RAP treatment resulted in effective depletion of HA-PfARK1. Western blot for β-actin was performed to serve as a loading control. **e**. HA-PfARK1-loxP parasites were synchronised and were harvested at Ring (R), Trophozoite (T) and Early Schizont (ES) or Late Schizont (LS) stages and Western blotting was performed using anti-HA or BiP antibodies. **f**. HA-PfARK1-loxP parasites were treated with DMSO or 250 nM RAP. Subsequently, RNA was isolated (~44 hpi, cycle1) and used for RT-qPCR analysis to assess the expression of ARK1/2/3 (SEM ± SE, sidak's multiple comparison, two-way ANOVA, $*P < 0.05$, $**P < 0.01$, ns = non-significant, $P$=0.0048 (ARK1), 0.042 (ARK2), and 0.9606 (ARK3), three independent biological replicates).

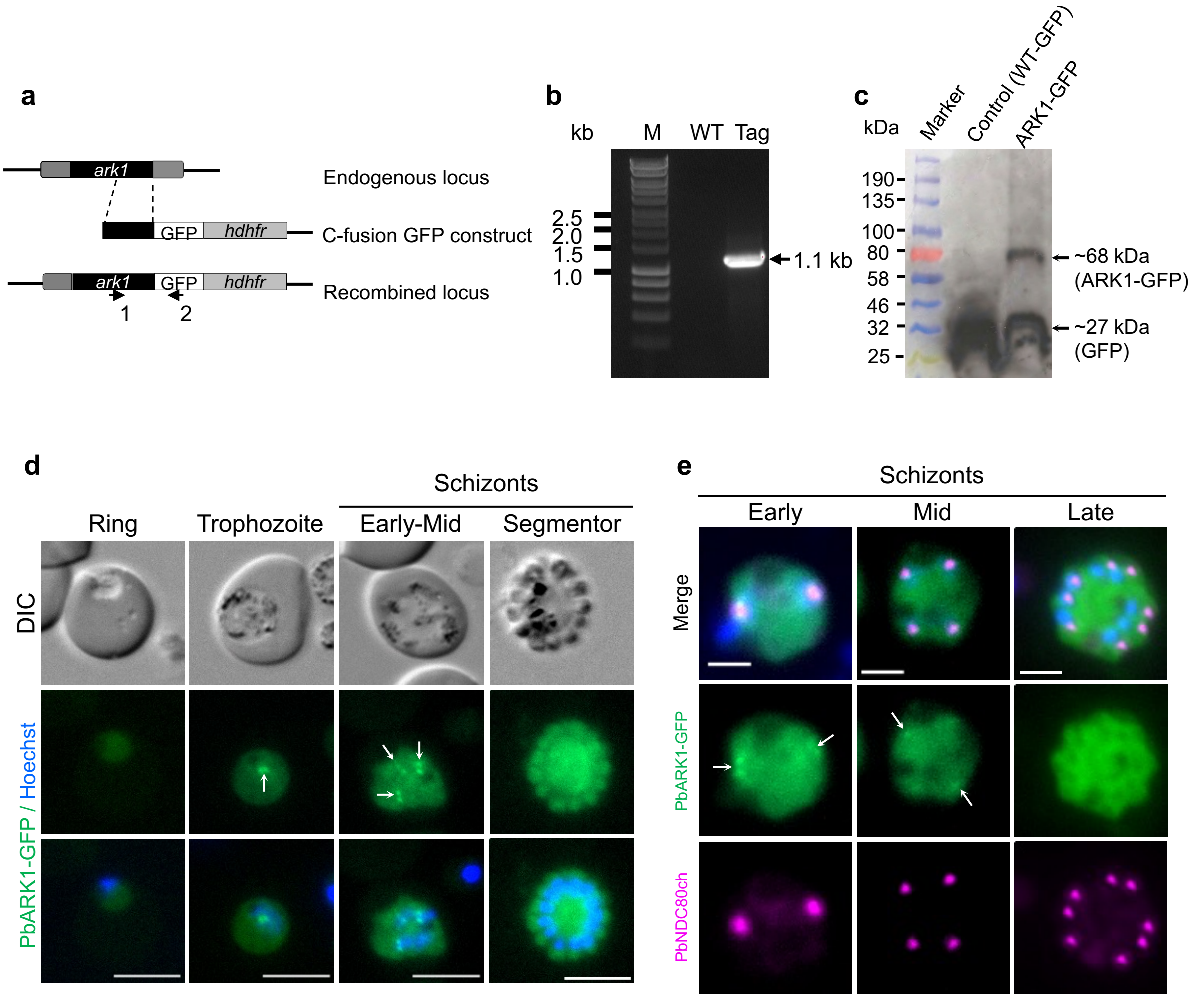


**Supplementary Figure 2. Generation of PbARK1-GFP parasites and analysis of subcellular location.**

**a.** Schematic representation of the endogenous *Pbark1* locus, the GFP-tagging construct, and the recombined *ark1* locus generated by single homologous recombination. Arrows 1 and 2 indicate the positions of PCR primers used to confirm successful integration of the construct. **b.** Diagnostic PCR of ark1 and WT parasites using primers IntArk1tg (Arrow 1) and ol492 (Arrow 2) to show the correct integration. Integration of the ark1 tagging construct gives a band of 1.1 kb. **c**. Western blot showing the expression of endogenous PbARK1-GFP detected by anti-GFP antibody. WT-GFP was used as control. The band in the control lane corresponds to GFP (~27 kDa). In the ARK1-GFP lane, the upper band corresponds to the full-length ARK1-GFP protein (~68 kDa), while the lower band likely represents a truncated protein containing GFP derived from ARK1-GFP. A detailed explanation of this data is provided in Supplementary Note 1. **d.** Live cell images showing the location of PbARK1-GFP during asexual blood stages of *P. berghei*. Representative of more than 50 images analysed in 3 different experiments. Scale bar = 5 μm. PbARK1-GFP foci are highlighted with white arrow. **e.** Live cell imaging showing the location of PbARK1-GFP (green) in relation to kinetochore marker PbNDC80-mCherry (magenta) during asexual blood stages of *P. berghei*. DNA is stained with Hoechst dye (blue). Merge: green, magenta, and blue images merged. Representative images of more than 20 cells with three biological replicates. Scale bar = 2 μm. PbARK1-GFP foci are highlighted with white arrow.

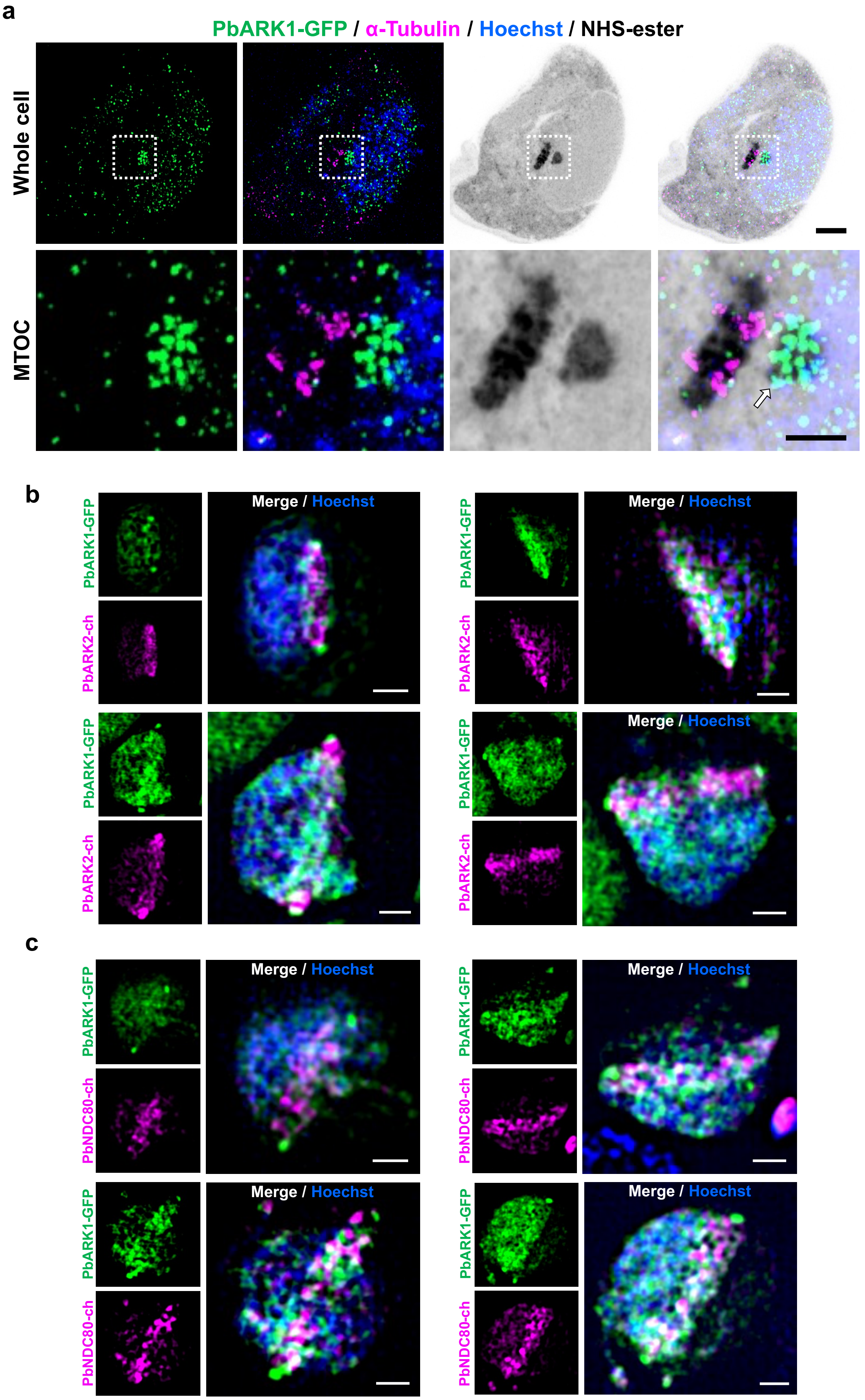

a
PbARK1-GFP / α-Tubulin / Hoechst / NHS-ester
Whole cell
MTOC
b
PbARK1-GFP
PbARK2-ch
Merge / Hoechst
c
PbNDC80-ch

**Supplementary Figure 3. U-ExM and 3D-SIM of PbARK1-GFP during *P. berghei* sexual male gametogony.**

**a.** U-ExM images of gametocytes at 1.5–2 min post activation (at the initial MTOC formation stage) showing location of PbARK1-GFP (green) and α- tubulin (magenta) detected with anti-GFP and anti-tubulin antibodies, respectively. NHS-ester and Hoechst were used to stain protein structures (grey) and DNA (blue), respectively. The top panels show the whole cell, and the bottom panels show magnified views of the MTOC indicated by white dashed squares. The PbARK1-GFP focus at the inner MTOC is indicated by arrows Scale bars = 5 µm (whole cell images) and 2 µm (spindle images). Representative images of more than 8 cells from two biological replicates. **b.** 3D-SIM images of fixed gametocytes at 1.5–2 min post activation (at the first spindle stage) showing the location of PbARK1-GFP and ARK2-mCherry. Representative image of more than 10 cells from more than two biological replicates. Scale bar = 1 µm. **c.** 3D-SIM images of fixed gametocytes at 1.5–2 min post activation (at the first spindle stage) showing the location of PbARK1-GFP and NDC80-mCherry. Representative image of more than 10 cells from more than two biological replicates. Scale bar = 1 µm.

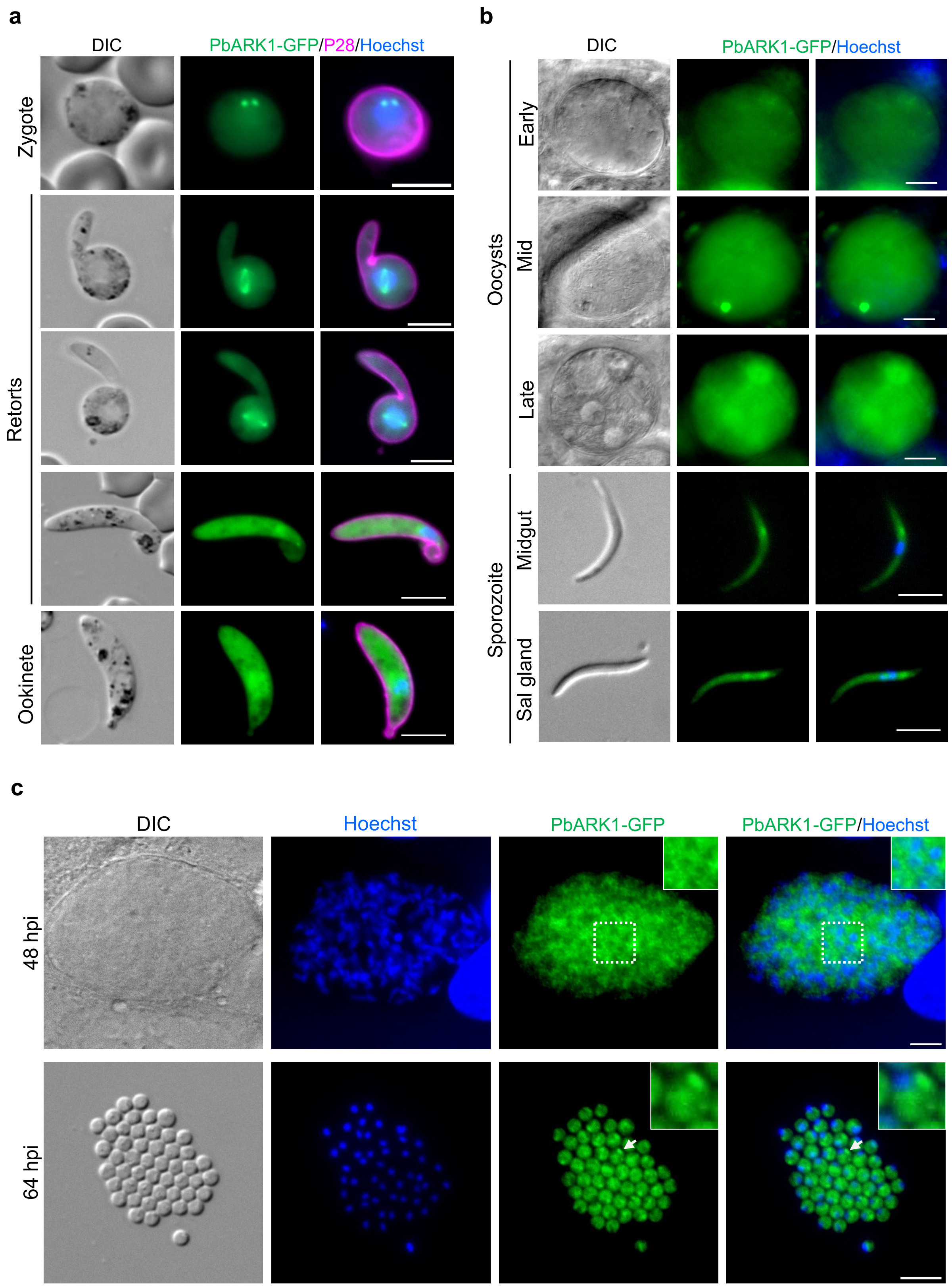


**Supplementary Figure 4. Realtime dynamics of PbARK1-GFP during meiosis and additional asexual developmental stages.**

**a.** Live-cell imaging showing PbARK1-GFP (green) location in zygote and during ookinete development, relative to the nuclear DNA (blue, Hoechst dye), and cy3-conjugated 13.1 antibody (magenta), which recognises P28 protein on the surface of zygotes and ookinetes. Representative of more than 50 images analysed in more than three different experiments. Scale bar = 5 µm. **b.** Live cell images showing the location of PbARK1-GFP during oocysts and sporozoite stages in mosquito. Representative of more than 50 images analysed in 3 different experiments. Scale bar = 5 µm. **c**. Live cell images showing the location of PbARK1-GFP in liver stages at 48 hours and 64 hours post infection (hpi). Insets show magnified views of the cells indicated by the dashed square (48 hpi) or arrow (64 hpi), respectively. Representative of more 20 images analysed in 3 different experiments. Scale bar = 5 µm. A detailed discussion of this data is provided in Supplementary Note 2.

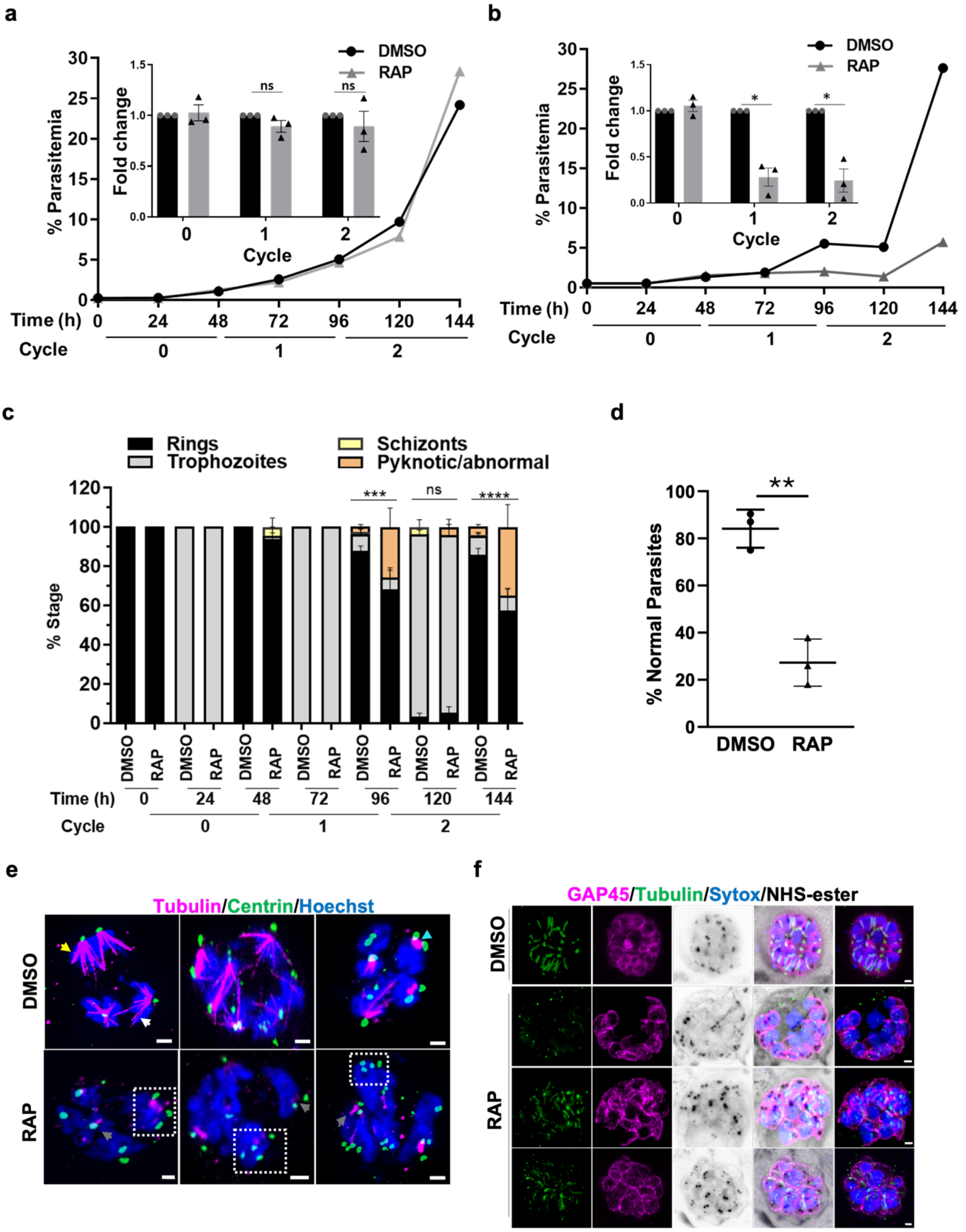


**Supplementary Figure 5. Effect of PfARK1 cKO on parasite development and division.**

**a** and **b**. The unmodified 1G5DC SkipFlox (a) or HA-PfARK1-loxP (b) parasites were synchronised and ring stage parasites and were either left untreated or treated with RAP as described for Figure 4a. Parasites were analysed at indicated time points by counting parasites by flow cytometry (a) or by microscopy of Giemsa-stained thin blood smears (b) and %

parasitaemia was determined [SEM ± SE, paired t test (a) cycle 1- ns, $P = 0.2036$, cycle 2- ns, $P = 0.5479$, (b) cycle 1- $*P = 0.0186$, cycle 2- $*P = 0.0267$, three independent biological replicates]. **c**. HA-PfARK1-loxP parasites were synchronized and ring stage parasites were treated with RAP as described for Figure 4a. The individual ring, trophozoites, and schizont stages were counted at the indicated time points from Giemsa-stained thin blood smears of parasite cultures and are represented as % of total number of parasites. In the case of RAP treated parasites, pyknotic morphology (Fig. 4b) was observed, and were also quantified separately (SEM ± SE, tukey's multiple comparison, two-way ANOVA, $***P = 0.0004$, $****P < 0.0001$, ns $P > 0.9999$, three independent biological replicates). **d**. This figure is the quantitation for Figure 4c. HA-PfARK1-loxP parasites were treated with DMSO or RAP and parasites were treated with E64 to prevent egress but allow complete maturation and division. Thin blood smears were made ~40-44 hpi in cycle 1 and nuclei were stained with DAPI. % parasites with normal segregated nuclei were counted (SEM ± SE, unpaired t test, $**P < 0.01$, n > 70 parasites, three independent biological replicates). **e**. HA-PfARK1-loxP parasites were treated with DMSO or RAP and U-ExM was performed on schizonts (cycle 1) as described for figure 4d (hemispindle - yellow arrow, mitotic spindles – cyan arrow, interpolar spindles – white arrow). RAP treatment resulted in a loss of spindles (grey arrow) in most parasites and unsegregated nuclei were observed and several CP clusters were observed which were often detached (square). **f.** Additional images showing the defects in cytokinesis as described in figure 4f. HA-PfARK1-loxP parasites were treated with DMSO or RAP and U-ExM was performed on schizonts (cycle 1). RAP treated parasites shows disorganised tubulin and aberrant GAP45 staining.

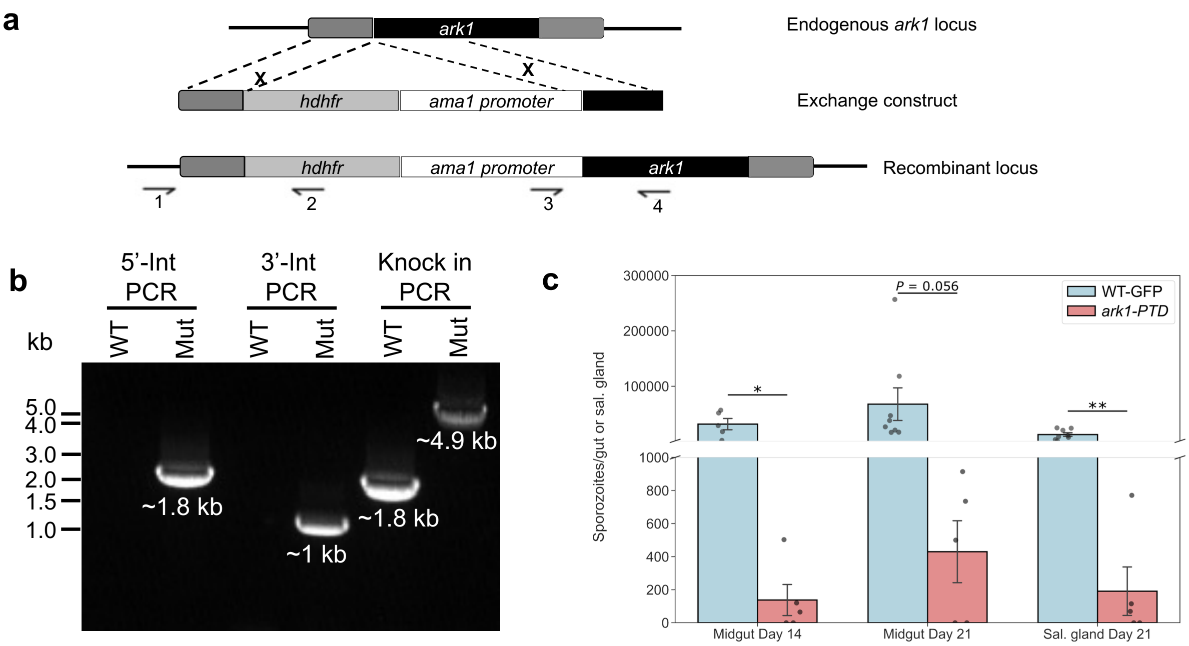


d

| Bite-back Experiment | | | | | |
|---|---|---|---|---|---|
| Exp | Parasite line | Blood Infection Day | Exp | Parasite line | Blood Infection Day |
| 1 | WT-GFP | 4 | 5 | WT-GFP | 4 |
| | *ark1-PTD* | No infection | | *ark1-PTD* | 14 |
| 2 | WT-GFP | 4 | 6 | WT-GFP | 4 |
| | *ark1-PTD* | No infection | | *ark1-PTD* | 10 |
| 3 | WT-GFP | 4 | 7 | WT-GFP | 4 |
| | *ark1-PTD* | 14 | | *ark1-PTD* | No infection |
| 4 | WT-GFP | 4 | 8 | WT-GFP | 4 |
| | *ark1-PTD* | No infection | | *ark1-PTD* | No infection |

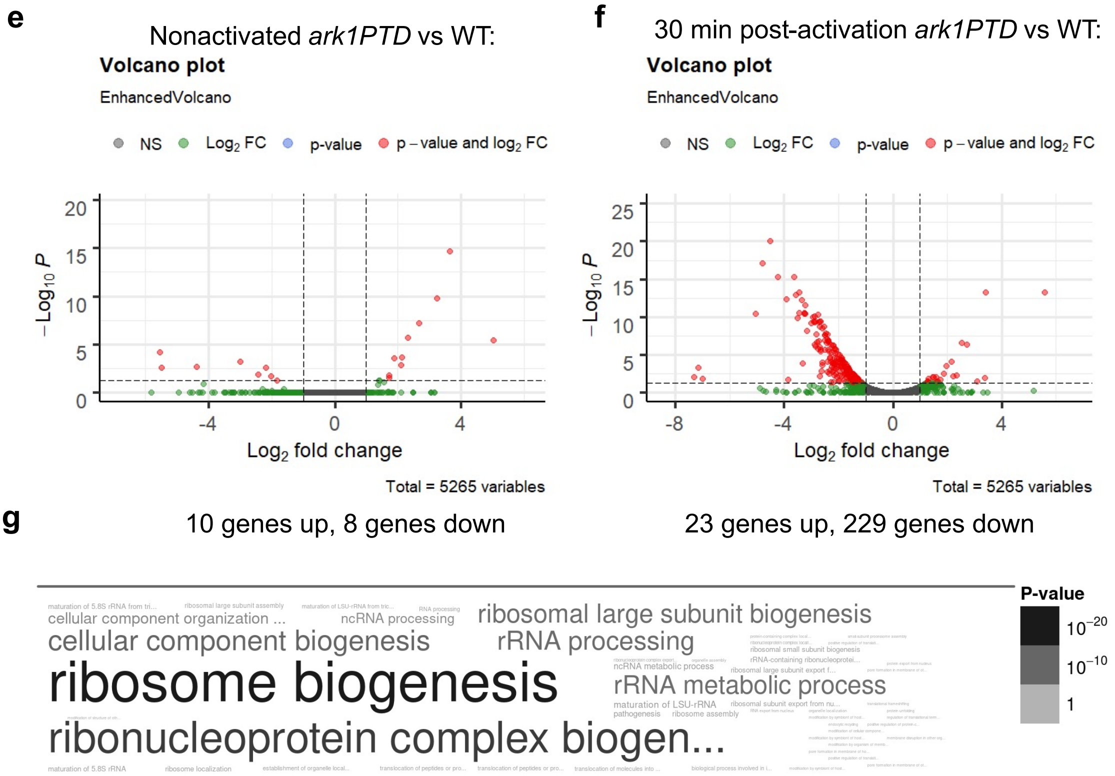

**Supplementary Figure 6. Generation of *Pbark1PTD* parasites and phenotypic analysis**.

**a.** Schematic representation of the promoter swap strategy to construct *Pbark1PTD* parasites (placing *ark1* under the control of the *ama1* promoter) by double homologous recombination. Arrows 1 and 2 indicate the primer positions used to confirm 5' integration and arrows 3 and 4 indicate the primers used to confirm 3' integration. **b.** Integration PCR of the promotor swap construct into the *ark1* locus. Primer 1 (IntPTD31_5) and primer 2 (5'-IntPTD) were used to confirm successful integration of the selectable marker, resulting in a band of 1.8 kb. Primer 3 (3'intPTama1) and primer 4 (IntPTD31_3) were used to determine the successful integration of the clag promoter, resulting in a band of 1kb. Primer 1 (IntPTD31_5) and primer 4 (IntPTD31_3) were used to confirm a complete knock-in of the construct with a band at 4.9 kb and the absence of a band at 1.8 kb. Representative image of more than 3 experiments. **c.** Total number of sporozoites per gut at day 14-, and 21 post-infections; and in salivary gland at day-21 post-infection in *ark1PTD* compared to WT-GFP parasites. Mean ± SEM. n ≥ 3 independent experiments**.** Statistical significance was determined using two-sided Welch's t-tests. **P* < 0.05, ***P* < 0.01. *P* = 0.037 (Midgut Day 14), 0.056 (Midgut Day 21), 0.006 (Salivary gland Day 21). **d.** Bite back experiment showing successful transmission of *ark1PTD* and WT-GFP parasites from mosquito to mouse with two different clones in three different experiments. **e.** Gene expression profile analysis using a volcano plot to highlight up- and downregulated genes in *ark1-PTD* compared to WT-GFP gametocytes (non-activated) with an adjusted *P*-value cutoff of 0.05. **f.** Gene expression profile analysis using a volcano plot to highlight up- and downregulated genes in *ark1-PTD* compared to WT-GFP gametocytes (activated for 30 min) with an adjusted *P*-value cutoff of 0.05. **g.** Gene ontology enrichment analysis for downregulated genes in *ark1-PTD* compared to WT-GFP gametocytes, activated for 30 min. A detailed discussion of this data is provided in Supplementary Note 3.

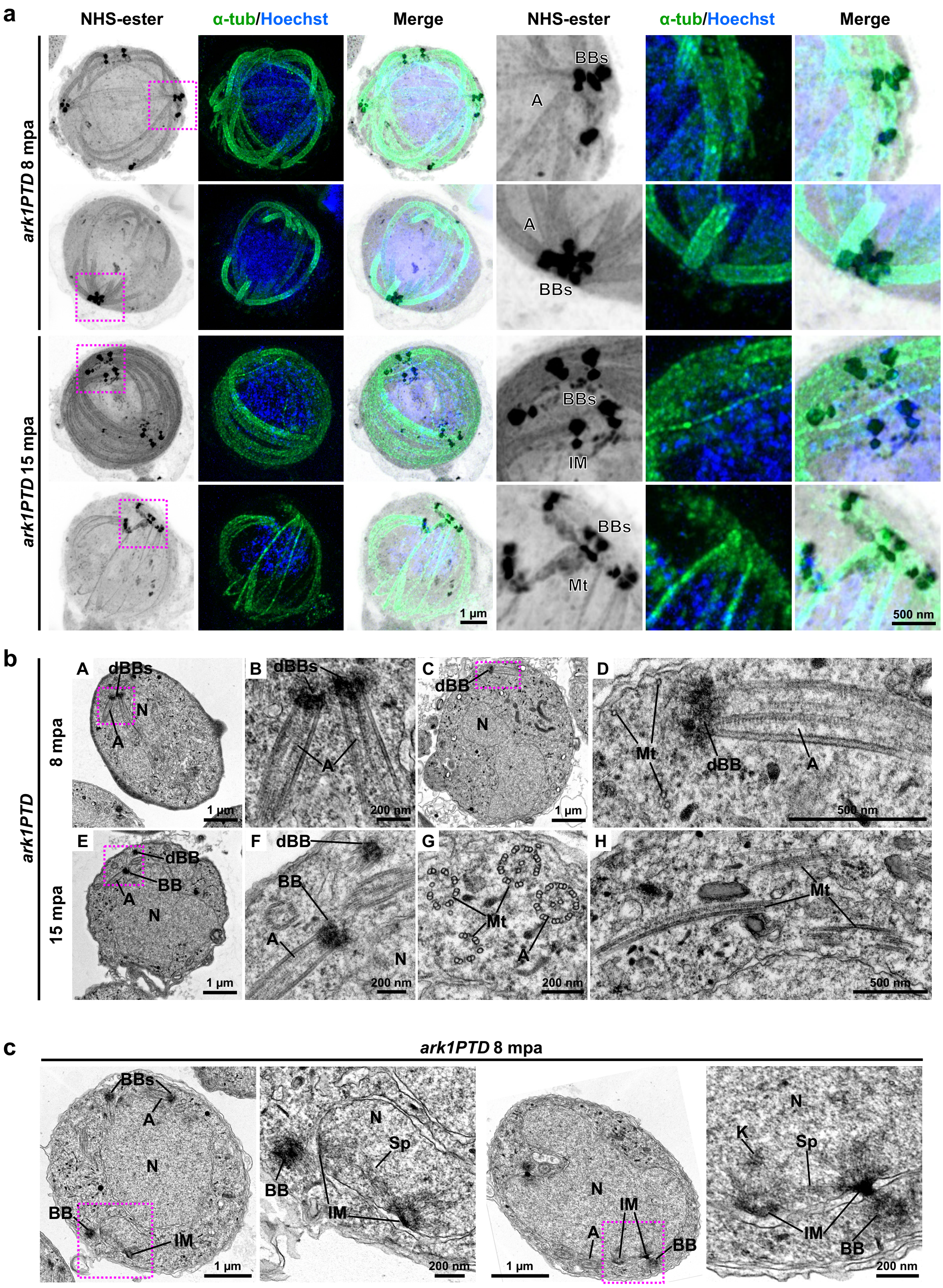


**Supplementary Figure 7. Additional expansion microscopy and transmission electron microscopy images of *ark1* gene-knockdown male gametocytes.**

**a.** *ark1* gene-knockdown (*ark1PTD*) male gametocytes at 8 min or 15 min post-activation (mpa). The three images on the left show maximum intensity projections of whole-cell z-stack images labelled with NHS-ester (grey), a-tubulin (green) and Hoechst (blue). The three images on the right show magnified views of the areas enclosed by the magenta squares. BBs: basal bodies (clumped), IM: inner MTOC, A: axoneme. Images are representative of more than 7 cells analysed across 3 independent experiments. The scale bars are scaled by an expansion factor of 4.954, calculated from a comparison of axoneme diameters obtained using expansion and transmission electron microscopy. **b.** Transmission electron microscopy images of *ark1* gene-knockdown (*ark1PTD*) male gametocytes at 8 min or 15 min post-activation. B, D and F are magnified views of the areas enclosed by the magenta squares in A, C and E, respectively. N: nucleus, BB: basal body, dBB: basal body that appeared detached from the nucleus, A: axoneme, Mt: microtubule. **c.** Additional transmission electron microscopy images revealing short spindle MTs spanning between two basal body MTOCs remained close together. IM: inner spindle MTOC, Sp: spindle microtubule, K: kinetochore. Images are representative of more than 50 cells analysed across 2 independent experiments.

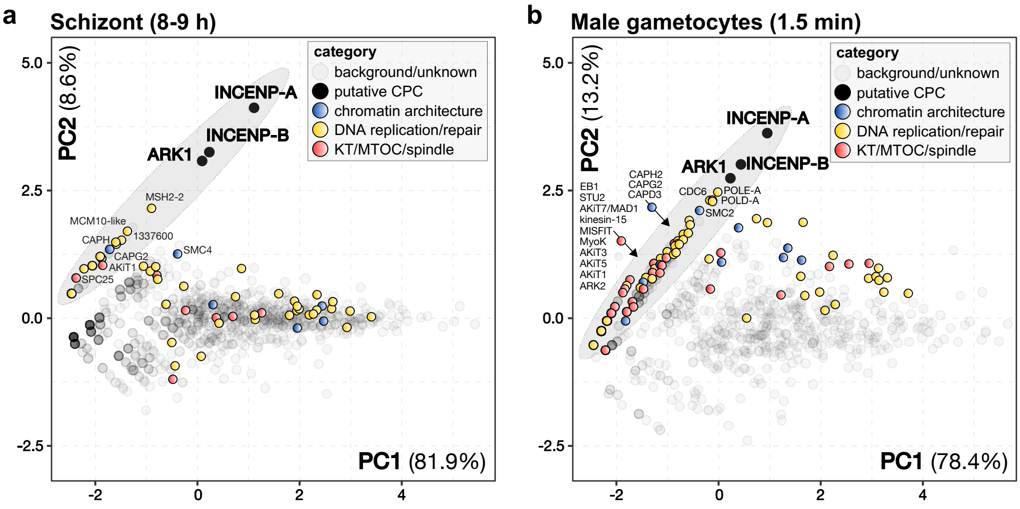


**c**

| Protein IDs | Description | gene_name | category | AA | Schizonts | | | | Male gametocytes | | | |
|---|---|---|---|---|---|---|---|---|---|---|---|---|
| | | | | | WT1 | WT2 | ARK1-1 | ARK1-2 | WT1 | WT2 | ARK1-1 | ARK1-2 |
| **PBANKA_1343200** | **inner centromere protein** | **INCENP-A** | **CPC** | **2305** | 0 | 0 | 60 | 44 | 0 | 0 | 76 | 41 |
| **PBANKA_1353900** | **inner centromere protein** | **INCENP-B** | **CPC** | **1126** | 0 | 0 | 21 | 21 | 0 | 0 | 42 | 28 |
| **PBANKA_0104100** | **aurora-related kinase ARK1** | **ARK1** | **CPC** | **353** | 0 | 0 | 13 | 15 | 0 | 0 | 24 | 13 |
| PBANKA_1011400 | DNA topoisomerase 2 | TOP2 | chromatin architecture | 1472 | 12 | 14 | 15 | 14 | 4 | 4 | 29 | 14 |
| PBANKA_1312000 | condensin-2 complex subunit G2 | CAPG2 | chromatin architecture | 1181 | 0 | 1 | 2 | 2 | 0 | 0 | 10 | 2 |
| PBANKA_0913000 | condensin-2 complex subunit D3 | CAPD3 | chromatin architecture | 1706 | 0 | 0 | 0 | 0 | 0 | 0 | 8 | 3 |
| PBANKA_0301800 | condensin-2 complex subunit H2 | CAPH2 | chromatin architecture | 775 | 0 | 0 | 0 | 0 | 0 | 0 | 6 | 2 |
| PBANKA_1337500 | protein STU2 | STU2 | kinetochore | 3198 | 3 | 1 | 5 | 2 | 0 | 0 | 14 | 6 |
| PBANKA_0612300 | apicomplexan kinetochore protein 7 | AKiT7/MAD1 | kinetochore | 832 | 2 | 1 | 0 | 2 | 0 | 0 | 10 | 2 |
| PBANKA_0612200 | apicomplexan kinetochore protein 3 | AKiT3 | kinetochore | 1251 | 0 | 0 | 0 | 0 | 0 | 0 | 8 | 0 |
| PBANKA_1243900 | apicomplexan kinetochore protein 5 | AKiT5 | kinetochore | 1522 | 1 | 1 | 1 | 2 | 0 | 0 | 6 | 0 |
| PBANKA_0621300 | apicomplexan kinetochore protein 1 | AKiT1 | kinetochore | 1737 | 1 | 1 | 2 | 2 | 0 | 0 | 4 | 3 |
| PBANKA_1358800 | kinetochore protein SPC25 | SPC25 | kinetochore | 216 | 0 | 0 | 2 | 0 | 0 | 0 | 3 | 0 |
| PBANKA_1115700 | kinetochore protein NDC80 | NDC80 | kinetochore | 589 | 0 | 0 | 0 | 0 | 0 | 0 | 3 | 0 |
| PBANKA_1442300 | kinetochore protein SPC24 | SPC24 | kinetochore | 819 | 0 | 0 | 0 | 0 | 0 | 0 | 3 | 0 |
| PBANKA_0522000 | apicomplexan kinetochore protein 6 | AKiT6 | kinetochore | 465 | 0 | 0 | 0 | 0 | 0 | 0 | 2 | 0 |
| PBANKA_0414300 | kinetochore protein NUF2 | NUF2 | kinetochore | 457 | 0 | 0 | 0 | 0 | 0 | 0 | 0 | 2 |
| PBANKA_0908500 | myosin K | MyoK | spindle | 1629 | 0 | 0 | 0 | 0 | 0 | 0 | 12 | 0 |
| PBANKA_1038200 | nuclear formin-like protein MISFIT | MISFIT | spindle | 1594 | 0 | 0 | 0 | 0 | 0 | 0 | 7 | 2 |
| PBANKA_0405600 | end-binding protein 1 | EB1 | spindle | 435 | 0 | 0 | 0 | 0 | 0 | 0 | 6 | 3 |
| PBANKA_1458800 | kinesin-15 | kinesin-15 | spindle | 1414 | 0 | 0 | 0 | 0 | 0 | 0 | 5 | 6 |
| PBANKA_0407400 | aurora-related kinase ARK2 | ARK2 | spindle | 1719 | 0 | 0 | 0 | 0 | 0 | 0 | 4 | 0 |
| PBANKA_0805900 | kinesin-8X | kinesin-8X | spindle | 1409 | 0 | 0 | 0 | 0 | 0 | 0 | 4 | 0 |
| PBANKA_0416100 | dynein heavy chain | DHC3 | MTOC | 5834 | 4 | 7 | 1 | 3 | 7 | 12 | 46 | 40 |
| PBANKA_0925400 | dynein heavy chain | DHC4A | MTOC | 5176 | 2 | 4 | 3 | 3 | 6 | 8 | 36 | 36 |
| PBANKA_0214000 | dynein heavy chain | DHC1 | MTOC | 5058 | 8 | 6 | 6 | 9 | 2 | 4 | 35 | 4 |
| PBANKA_0507300 | dynein heavy chain | DHC4B | MTOC | 5309 | 0 | 4 | 1 | 0 | 4 | 6 | 29 | 31 |
| PBANKA_0801500 | conserved Plasmodium protein | CCDC63 | MTOC | 745 | 0 | 0 | 0 | 0 | 0 | 2 | 14 | 9 |
| PBANKA_0202700 | kinesin-8B | kinesin-8B | MTOC | 1460 | 0 | 0 | 0 | 0 | 2 | 10 | 13 | 10 |
| PBANKA_1361500 | WD repeat-containing protein 16 | CFAP52 | MTOC | 723 | 0 | 2 | 0 | 0 | 0 | 2 | 12 | 10 |
| PBANKA_0102700 | conserved Plasmodium protein | CFAP157 | MTOC | 372 | 0 | 0 | 0 | 0 | 0 | 0 | 8 | 4 |
| PBANKA_1430900 | SUN domain-containing protein | SUN1 | MTOC | 872 | 0 | 0 | 0 | 0 | 0 | 0 | 6 | 0 |
| PBANKA_1018300 | dynein intermediate chain | DNAI2 | MTOC | 592 | 2 | 5 | 2 | 1 | 0 | 3 | 5 | 9 |
| PBANKA_0104200 | calcium-binding protein | 104200 | MTOC | 157 | 0 | 0 | 0 | 0 | 0 | 0 | 4 | 3 |
| PBANKA_1035700 | WD repeat-containing protein 65 | CFAP57/WDR65 | MTOC | 1440 | 0 | 0 | 0 | 0 | 0 | 0 | 4 | 0 |
| PBANKA_0507800 | conserved protein | CFAP100 | MTOC | 492 | 0 | 0 | 0 | 0 | 0 | 0 | 3 | 2 |
| PBANKA_1125300 | centrosomal protein CEP135 | CFAP58/CEP135 | MTOC | 906 | 0 | 0 | 0 | 0 | 0 | 0 | 3 | 0 |
| PBANKA_1312800 | CLASP domain-containing protein | CLASP | MTOC | 859 | 0 | 0 | 0 | 0 | 0 | 0 | 3 | 0 |
| PBANKA_1446000 | conserved Plasmodium protein | CEP104 | MTOC | 916 | 0 | 0 | 0 | 0 | 0 | 0 | 3 | 0 |
| PBANKA_1109600 | conserved Plasmodium protein | RSP9 | MTOC | 672 | 2 | 2 | 2 | 2 | 0 | 0 | 3 | 2 |
| PBANKA_0934700 | conserved protein | CFAP73/CCDC42 | MTOC | 362 | 0 | 0 | 0 | 0 | 0 | 0 | 2 | 3 |
| PBANKA_0504300 | flagellar outer arm dynein-associated protein | DYNLRB1 | MTOC | 101 | 0 | 0 | 0 | 0 | 0 | 0 | 2 | 2 |
| PBANKA_1133600 | dynein light chain | 1133600 | MTOC | 121 | 0 | 0 | 0 | 0 | 0 | 0 | 2 | 2 |
| PBANKA_0415800 | WD repeat-containing protein 66 | CFAP251 | MTOC | 1111 | 0 | 0 | 0 | 0 | 0 | 0 | 2 | 0 |
| PBANKA_1443000 | NIMA related kinase 1 | NEK1 | MTOC | 915 | 0 | 0 | 0 | 0 | 0 | 0 | 2 | 0 |
| PBANKA_0708100 | dynein light chain 1 | DLC1 | MTOC | 198 | 0 | 0 | 0 | 0 | 0 | 0 | 0 | 2 |
| PBANKA_1108000 | CFA20 domain-containing protein | CFAP20 | MTOC | 364 | 0 | 0 | 0 | 0 | 3 | 4 | 0 | 3 |
| PBANKA_1454600 | nucleoporin NUP269 | NUP269 | MTOC | 2260 | 0 | 0 | 0 | 0 | 0 | 0 | 7 | 0 |
| PBANKA_0100800 | zinc finger protein | 100800 | other | 1382 | 2 | 3 | 5 | 4 | 0 | 0 | 19 | 7 |
| PBANKA_0818500 | conserved protein | 818500 | other | 665 | 2 | 2 | 3 | 1 | 0 | 0 | 15 | 12 |
| PBANKA_1417900 | conserved protein | 1417900 | other | 1052 | 1 | 3 | 1 | 2 | 0 | 2 | 14 | 5 |
| PBANKA_0314200 | calcium-dependent protein kinase 1 | CDPK1 | other | 523 | 2 | 1 | 2 | 2 | 3 | 2 | 13 | 9 |
| PBANKA_1426500 | cullin-1 | CUL1 | other | 844 | 1 | 1 | 1 | 2 | 0 | 0 | 12 | 3 |
| PBANKA_1003400 | amino acid decarboxylase AAD | UIS14 | other | 2262 | 1 | 1 | 0 | 2 | 0 | 0 | 11 | 0 |
| PBANKA_1320100 | conserved Plasmodium protein | UIS25 | other | 789 | 0 | 0 | 0 | 0 | 0 | 2 | 10 | 5 |
| PBANKA_0933700 | mitogen-activated protein kinase 2 | MAPK2 | other | 523 | 0 | 0 | 0 | 0 | 0 | 2 | 7 | 7 |
| PBANKA_1129100 | DNA polymerase epsilon catalytic subunit A | POLE-A | DNA replication & repair | 2826 | 7 | 7 | 9 | 10 | 0 | 0 | 41 | 17 |
| PBANKA_1034100 | FANCJ-like helicase | FANCJ | DNA replication & repair | 1069 | 3 | 4 | 10 | 7 | 0 | 0 | 31 | 13 |
| PBANKA_0613200 | DNA polymerase alpha catalytic subunit A | POLA-A | DNA replication & repair | 1709 | 4 | 5 | 7 | 6 | 0 | 3 | 30 | 17 |
| PBANKA_1442200 | protein SOC2 | SOC2 | DNA replication & repair | 5372 | 1 | 0 | 5 | 3 | 0 | 0 | 25 | 7 |
| PBANKA_0712100 | DNA helicase | 712100 | DNA replication & repair | 1018 | 2 | 3 | 5 | 5 | 0 | 0 | 24 | 18 |
| PBANKA_0804300 | DNA mismatch repair protein MSH2 | MSH2-2 | DNA replication & repair | 853 | 1 | 1 | 7 | 6 | 4 | 8 | 24 | 15 |
| PBANKA_1302600 | DNA primase small subunit | PRIM1 | DNA replication & repair | 447 | 2 | 2 | 3 | 2 | 0 | 2 | 15 | 7 |
| PBANKA_0307100 | conserved Plasmodium protein | Timeless/Tof1 | DNA replication & repair | 2137 | 2 | 2 | 5 | 3 | 0 | 0 | 14 | 10 |
| PBANKA_1358300 | conserved Plasmodium protein | MSH2 | DNA replication & repair | 1419 | 0 | 1 | 1 | 2 | 0 | 0 | 12 | 4 |
| PBANKA_1337600 | BRCT domain-containing protein | 1337600 | DNA replication & repair | 1265 | 1 | 0 | 3 | 3 | 0 | 0 | 11 | 5 |
| PBANKA_0812000 | DNA primase large subunit | PRIM2 | DNA replication & repair | 537 | 2 | 0 | 3 | 4 | 0 | 0 | 10 | 6 |
| PBANKA_1230200 | cdc2-related protein kinase 5 | CRK5 | DNA replication & repair | 712 | 0 | 0 | 0 | 0 | 0 | 0 | 8 | 0 |
| PBANKA_1326900 | DNA polymerase alpha subunit B | POLA-B | DNA replication & repair | 541 | 0 | 3 | 5 | 2 | 0 | 0 | 8 | 8 |
| PBANKA_1006100 | flap endonuclease 1 | FEN1 | DNA replication & repair | 479 | 2 | 1 | 1 | 2 | 2 | 3 | 8 | 4 |
| PBANKA_1356400 | DNA repair protein RAD5 | RAD5 | DNA replication & repair | 1433 | 0 | 0 | 0 | 0 | 0 | 0 | 7 | 3 |
| PBANKA_0306500 | ARMT1-like domain-containing protein | ARMT1 | DNA replication & repair | 454 | 5 | 3 | 3 | 0 | 0 | 0 | 7 | 2 |
| PBANKA_1133800 | DNA helicase | 1133800 | DNA replication & repair | 867 | 0 | 0 | 0 | 0 | 0 | 0 | 6 | 3 |
| PBANKA_1349400 | DNA replication complex GINS protein | PSF2 | DNA replication & repair | 502 | 0 | 0 | 0 | 0 | 0 | 0 | 5 | 4 |
| PBANKA_1448900 | DNA polymerase epsilon subunit B | POLE-B | DNA replication & repair | 623 | 1 | 0 | 1 | 2 | 0 | 0 | 5 | 2 |
| PBANKA_1344300 | chromatin assembly factor 1 subunit B | CAF1B | DNA replication & repair | 513 | 0 | 0 | 0 | 0 | 0 | 0 | 4 | 3 |
| PBANKA_1102900 | cell division control protein 6 | CDC6 | DNA replication & repair | 869 | 1 | 2 | 3 | 2 | 0 | 0 | 24 | 15 |
| PBANKA_1356300 | CDT1-like protein | CDT1 | DNA replication & repair | 955 | 4 | 5 | 0 | 5 | 0 | 0 | 19 | 8 |
| PBANKA_0312500 | origin recognition complex subunit 5 | ORC5 | DNA replication & repair | 815 | 0 | 0 | 3 | 0 | 0 | 0 | 16 | 7 |
| PBANKA_1348800 | origin recognition complex subunit 4 | ORC4 | DNA replication & repair | 885 | 2 | 0 | 3 | 4 | 0 | 0 | 12 | 8 |
| PBANKA_0513900 | ORC3 domain-containing protein | ORC3 | DNA replication & repair | 1346 | 0 | 0 | 2 | 1 | 0 | 0 | 11 | 8 |
| PBANKA_1336900 | conserved Plasmodium protein | MCM10-like | DNA replication & repair | 508 | 0 | 0 | 3 | 4 | 0 | 0 | 11 | 7 |
| PBANKA_0602000 | origin recognition complex subunit 1 | ORC1 | DNA replication & repair | 1038 | 0 | 0 | 0 | 0 | 0 | 0 | 11 | 5 |
| PBANKA_0803000 | origin recognition complex subunit 2 | ORC2 | DNA replication & repair | 781 | 1 | 1 | 4 | 2 | 0 | 0 | 10 | 4 |
| PBANKA_0202500 | nucleoside triphosphate hydrolase | RFC-like | DNA replication & repair | 1043 | 0 | 0 | 2 | 1 | 0 | 0 | 8 | 2 |
| PBANKA_0303300 | conserved Plasmodium protein | CDC45 | DNA replication & repair | 619 | 1 | 1 | 3 | 1 | 0 | 0 | 8 | 8 |

**Supplementary Figure 8. ARK1 interactomics in schizonts and male gametocytes.** Principal Component Analysis (PCA) of duplicate **a.** schizont 8-9 h and **b.** male gametocytes 1.5 min lysates for GFP-trap of ARK1-GFP vs. WT-GFP. **c.** Table with unique peptide values for ARK1, INCENP-A/-B and other associated proteins in both male gametocytes and schizonts. Values can be found in Supplementary Data 2**.**

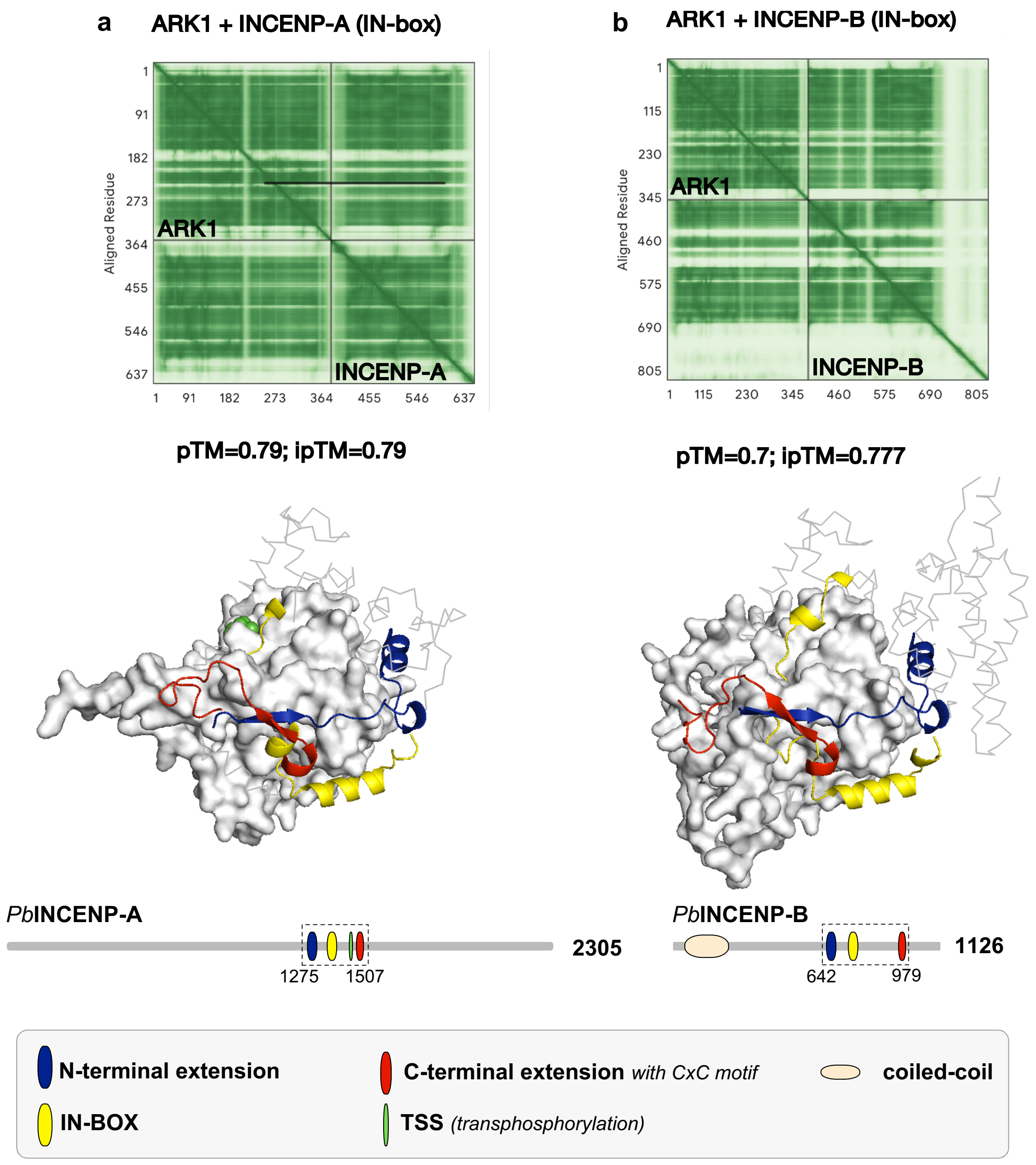


**Supplementary Figure 9. Aligned Error (AE) plot for AF3 predictions for ARK1-INCENP-A and ARK1-INCENP-B predictions.**

**a.** ARK1-INCENP-A prediction. **b.** ARK1-INCENP-B prediction. (i)pTM scores > 0.7 suggest a robustly predicted interaction. 3D protein structures and protein cartoons are the same as Figure 7b.

#### Supplementary Notes

##### Supplementary Note 1. Generation of PbARK1-GFP line.

PbARK1-GFP was integrated by single crossover homologous recombination at the 3′ end of the endogenous *Pbark1* locus (Supplementary Fig. 2a). Correct integration of the tagging construct was confirmed by PCR analysis of genomic DNA using locus-specific diagnostic primers (Supplementary Fig. 2b). Western blot analysis detected a PbARK1-GFP fusion protein of the expected size (~68 kDa) (Supplementary Fig. 2c). The PbARK1-GFP parasites were viable and completed the full life cycle with no detectable phenotypic defects resulting from the GFP tagging.

##### Supplementary Note 2. Localisation of PbARK1-GFP during other mosquito stages and liver stages.

We took advantage of the experimental accessibility of *P. berghei* to investigate the localisation of PbARK1-GFP during meiotic and other asexual stages in the mosquito. Live cell imaging revealed a diffuse distribution of PbARK1-GFP in the cytoplasm in zygote and throughout ookinete development (Supplementary Fig. 4a). Additionally, two closely associated focal points of PbARK1-GFP were detected in the zygote approximately 2 hours post-fertilization. These focal points subsequently moved apart, localising to opposite poles of the nucleus during ookinete transformation, before disappearing in the late and mature stages (Supplementary Fig. 4a). Mature ookinetes traverse the mosquito midgut and form oocysts on its outer surface. We examined the localisation of PbARK1-GFP during oocyst development and observed a diffuse distribution throughout the oocyst (Supplementary Fig. 4b). Similarly, sporozoites developing within oocysts, as well as those collected from the salivary glands, also exhibited a diffuse localisation pattern of PbARK1-GFP (Supplementary Fig. 4b). Next, we infected HepG2 cells with these sporozoites to study PbARK1-GFP localisation during the liver stage of infection. Live cell imaging revealed a diffuse distribution of PbARK1-GFP in early liver stage parasites (48 hours post-infection), with a slight enrichment of PbARK1-GFP adjacent to the nucleus observed in the late stages (64 hours post-infection) (Supplementary Fig. 4c).

##### Supplementary Note 3. Gene expression profile analysis.

To investigate the effect of *Pbark1* downregulation at the genome wide level, we performed RNA-seq analysis in triplicate, starting with total RNA extracted from purified *ark1PTD* and WT-GFP gametocytes at two time points: before activation (non-activated, 0 min) and at 30 min post-activation (mpa). Differences in gene expression between the correspondent samples were detected. In non-activated gametocytes, only a small number of differentially expressed genes were identified in the *ark1PTD* sample, with 10 genes up- and 8 genes downregulated, respectively (Supplementary Fig. 6e and Supplementary Data 1). Those genes included several fam a and b proteins known to be highly variable during life cycle progression due to their involvement in host immune response modulation[3]. They were therefore not considered as significant**.** At 30 mpa, 23 genes were upregulated including several conserved *Plasmodium* proteins with unknown function as well as PBANKA_1206900, a tubulin beta chain and putatively critical during gametocyte formation[4]. A significantly higher number of genes (229) were downregulated (Supplementary Fig. 6f and Supplementary Data 1). These genes included several protein kinases, e.g. PBANKA_1228800 and calcium-dependent protein kinase 5 (CDPK5 – PBANKA_1351500), as well as genes involved in lipid biosynthesis including PBANKA_0201300, a putative lysophospholipase, PBANKA_1127000, and a choline/ethanolaminephosphotransferase. Also, down-regulated were myosin essential light chain (ELC) and skeleton-binding protein 1 (PBANKA_1101300), as well as several enzymes known to be critical for sexual differentiation. Gene ontology (GO) analysis also revealed that several genes detected as downregulated were associated with ribosome biogenesis (Supplementary Fig. 6g), including ribosome biogenesis protein RPF2 putative (PBANKA_0512500) and ribosome biogenesis protein MRT4 putative (PBANKA_1143500). While an alternative interpretation may involve regulatory turnover post-fertilisation, this broad downregulation is consistent with the observed developmental arrest in *ark1PTD* parasites.
